# Treatment with furosemide indirectly increases inhibitory transmission in the developing hippocampus

**DOI:** 10.1101/2023.07.11.548438

**Authors:** C. Peerboom, T. Wijne, C.J. Wierenga

**Affiliations:** Cell Biology, Neurobiology and Biophysics, Biology department, Utrecht University, 3584 CH, Utrecht, the Netherlands; Donders Institute for Brain, Cognition and Behaviour, Radboud University, 6525 AJ, Nijmegen, the Netherlands

**Author notes:** Corresponding author: Corette J. Wierenga. ***Author contributions:*** C.P., and C.J.W. designed research; C.P and T.W., performed research and analyzed data; C.P., and C.J.W. wrote the paper.

## Abstract

During the first two postnatal weeks intraneuronal chloride concentrations in rodents gradually decrease, causing a shift from depolarizing to hyperpolarizing γ-aminobutyric acid (GABA) responses. GABAergic depolarization in the immature brain is crucial for the formation and maturation of excitatory synapses, but when GABAergic signaling becomes inhibitory it no longer promotes synapse formation. Here we examined the role of chloride transporters in developing postnatal hippocampal neurons using furosemide, an inhibitor of the chloride importer NKCC1 and chloride exporter KCC2 with reported anticonvulsant effects. We treated organotypic hippocampal cultures made from 6 to 7-day old mice with 200 μM furosemide from DIV1 to DIV8. Using perforated patch clamp recordings we observed that the GABA reversal potential was depolarized after acute furosemide application, but after a week of furosemide treatment the GABA reversal potential but was more hyperpolarized compared to control. Expression levels of the chloride cotransporters were unaffected after one week furosemide treatment. This suggests that furosemide inhibited KCC2 acutely, while prolonged treatment resulted in (additional) inhibition of NKCC1, but we cannot exclude changes in HCO_3_^-^. We assessed the effects of accelerating the GABA shift by furosemide treatment on inhibitory synapses onto CA1 pyramidal cells. Directly after cessation of furosemide treatment at DIV9, inhibitory synapses were not affected. However at DIV21, two weeks after ending the treatment, we found that the frequency of inhibitory currents was increased, and VGAT puncta density in *stratum Radiatum* was increased. In addition, cell capacitance of CA1 pyramidal neurons was reduced in furosemide-treated slices at DIV21 in an activity-dependent manner. Our results suggest that furosemide treatment indirectly promoted inhibitory transmission, possibly by increasing activity-independent GABA release.

## Introduction

γ-Aminobutyric acid (GABA) is the main inhibitory, hyperpolarizing neurotransmitter in the adult brain, but during early development GABA actually depolarizes neurons. Ionotropic GABA_A_ receptors primarily conduct chloride (∼70%), but also HCO_3_^-^ ions (∼30%) (Kaila et al., 2014). While HCO_3_^-^ levels remain constant, intracellular chloride levels decrease during brain development (Kaila et al., 2014). This developmental decrease in chloride causes the reversal potential (E_GABA_) to gradually drop below resting membrane potential. As a result, the GABAergic driving force shifts from depolarizing in immature neurons to hyperpolarizing in the mature brain (Rivera et al., 1999; Hübner et al., 2001; Tyzio et al., 2007; Romo-Parra et al., 2008; Kirmse et al., 2015; Tsukahara et al., 2015; Sulis Sato et al., 2017; Murata and Colonnese, 2020). The decrease in chloride levels is the direct result of an increased function of the chloride exporter KCC2 (K-Cl cotransporter 2) relative to the chloride importer NKCC1 (Na-K-2Cl cotransporter 1) (Rivera et al., 1999; Gulyás et al., 2001; Yamada et al., 2004; Dzhala et al., 2005; Otsu et al., 2020). In humans, the shift in chloride transporter expression occurs during the first year after birth (Dzhala et al., 2005; Sedmak et al., 2016). In rodents, the GABA shifts during the first two postnatal weeks (Rivera et al., 1999; Stein et al., 2004; Tyzio et al., 2007; Romo-Parra et al., 2008; Kirmse et al., 2015; Sulis Sato et al., 2017; Murata and Colonnese, 2020).

Depolarizing GABA signaling plays a critical role in the formation and maturation of excitatory synapses in the developing brain (Leinekugel et al., 1997; Akerman and Cline, 2006; Wang and Kriegstein, 2008, 2011; Chancey et al., 2013; van Rheede et al., 2015; Oh et al., 2016). However, after the first postnatal week, GABA becomes inhibitory and GABA signaling no longer promotes synapse formation (Salmon et al., 2020; Peerboom et al., 2023). Here we examined the role of chloride transporters in the development of inhibitory synapses in organotypic hippocampal cultures using furosemide, a well-known inhibitor of chloride transport. Treatment with furosemide has been employed previously to block KCC2 and elevate chloride concentrations in (slice) cultures (Thompson and Gahwiler, 1989; Jarolimek et al., 1999; Deeb et al., 2013; Wright et al., 2017), as furosemide has a preference for inhibiting KCC2 over NKCC1 (IC_50_ ∼30 μM for rat NKCC1 and IC_50_∼10 μM for rat KCC2, expressed in Xenopus oocytes (Orlov et al., 2015)). In addition, furosemide may interfere with α6- and α4-subunit containing GABA_A_ receptors (Thompson and Gahwiler, 1989; Pearce, 1993; Korpi et al., 1995), and with the regulation of neuronal HCO_3_^-^ levels (Halligan et al., 1991; Temperini et al., 2009; Ruusuvuori and Kaila, 2014; Uwera et al., 2015). Moreover, furosemide was shown to inhibit activity-induced swelling of epileptic brain tissue (Hochman et al., 1995; Gutschmidt et al., 1999; Hochman, 2012).

We examined the effects of 200 μM furosemide in organotypic hippocampal cultures. We observed that acute furosemide application depolarized GABA reversal potential, in line with a preferential inhibition of KCC2. In contrast, furosemide treatment from DIV1 to DIV8 resulted in hyperpolarization of the GABA reversal potential. Expression levels of the chloride-cotransporters were unaffected after furosemide-treatment. This suggests that sustained application of furosemide resulted in inhibition of NKCC1, although we cannot exclude changes in HCO_3_^-^ levels. We assessed the consequences of the accelerated GABA shift during furosemide treatment on inhibitory transmission. We found that inhibitory synapses were not affected directly after furosemide treatment at DIV9. However at DIV21, when E_GABA_ had normalized, inhibitory currents were strongly increased in furosemide-treated slices, possibly via an increased number of inhibitory synapses in the *stratum Radiatum*. In addition, we found evidence for shrinkage of CA1 pyramidal neurons in furosemide-treated slices at DIV21. Our results suggest that furosemide may indirectly promote inhibitory transmission.

## Methods

### Animals

All animal experiments were performed in compliance with the guidelines for the welfare of experimental animals issued by the Federal Government of the Netherlands and were approved by the Dutch Central Committee Animal experiments (CCD), project AVD1080020173847 and project AVD1150020184927. In this study we used male and female transgenic mice: GAD65-GFP mice (López-Bendito et al., 2004) (bred as a heterozygous line in a C57BL/6JRj background), C57BL/6JRj littermates, and SuperClomeleon (SClm) mice. The SClm mice (Herstel et al., 2022) are SuperClomeleon^lox/-^ mice (Rahmati et al., 2021), a gift from Kevin Staley (Massachusetts General Hospital, Boston, MA) crossed with CamKIIα^Cre/-^ mice (Tsien et al., 1996; Casanova et al., 2001a). SClm mice express the chloride sensor SClm in up to 70% of the pyramidal neurons in the hippocampus (Casanova et al., 2001b; Wang et al., 2013). GAD65-GFP mice express GFP in ∼20% of GABAergic interneurons in the CA1 region of the hippocampus (Wierenga et al., 2010). As we did not detect any differences between slices from male and female mice as well as between GAD65-GFP mice and C57BL/6JRj littermates, all data were pooled.

### Organotypic hippocampal culture preparation and furosemide-treatment

Organotypic hippocampal cultures were made from P6-7 mice as described before (Hu et al., 2019; Herstel et al., 2022), based on the *Stoppini method* (Stoppini et al., 1991). Mice were decapitated and their brain was rapidly placed in ice-cold Grey’s Balanced Salt Solution (GBSS; containing (in mM): 137 NaCl, 5 KCl, 1.5 CaCl_2_, 1 MgCl_2_, 0.3 MgSO_4_, 0.2 KH_2_PO_4_ and 0.85 Na_2_HPO_4_) with 25 mM glucose, 12.5 mM HEPES and 1 mM kynurenic acid (pH set at 7.2, osmolarity set at ∼320 mOsm, sterile filtered). 400 μm thick transverse hippocampal slices were cut with a McIlwain tissue chopper. Slices were placed on Millicell membrane inserts (Millipore) in wells containing 1 ml culture medium (consisting of 48% MEM, 25% HBSS, 25% horse serum, 25 mM glucose, and 12.5 mM HEPES, with an osmolarity of ∼325 mOsm and a pH of 7.3 – 7.4). Slices were stored in an incubator (35°C with 5% CO_2_). At DIV1 culture medium was replaced by culture medium supplemented with 0.1% dimethyl sulfoxide (DMSO, Sigma-Aldrich) or 200 μM furosemide (furosemide, Merck, in 0.1 % DMSO). In addition, a small drop of medium supplemented with DMSO or furosemide was carefully placed on top of the slices. In this way, cultures were treated 3 times per week until DIV8. From DIV8 onwards, cultures received normal culture medium 3 times per week. Experiments were performed at day in vitro 1-3 (DIV2), 8-10 (DIV9) or 20-22 (DIV21). Please note that perforated patch experiments at DIV9 were performed 1 to 55 hr after cessation of the furosemide treatment. E_GABA_ did not correlate with hours after treatment and was not affected by addition of furosemide to the ACSF during the perforated patch recording.

### Electrophysiology and analysis

Organotypic cultures were transferred to a recording chamber and continuously perfused with carbonated (95% O_2_, 5% CO_2_) artificial cerebrospinal fluid (ACSF, in mM: 126 NaCl, 3 KCl, 2.5 CaCl_2_, 1.3 MgCl_2_, 26 NaHCO_3_, 1.25 NaH_2_PO_4_, 20 glucose; with an osmolarity of ∼310 mOsm) at a rate of approximately 1 ml/min. Bath temperature was maintained at 29-32°C. Recordings were acquired using an MultiClamp 700B amplifier (Molecular Devices), filtered with a 3 kHz Bessel filter and stored using pClamp10 software. For acute application, ACSF containing 0.1% DMSO or 200 μM furosemide (dissolved in 0.1% DMSO) was bath applied for 5 minutes.

Perforated patch clamp recordings were performed in CA1 pyramidal neurons. Recording pipettes with resistances of 2-4 MΩ were pulled from borosilicate glass capillaries (World Precision Instruments). The pipette tip was filled with gramicidin-free KCl solution (140 mM KCl and 10 mM HEPES, pH adjusted to 7.2 with KOH, and osmolarity 285 mOsm/l) and then backfilled with solution containing gramicidin (60 µg/ml, Sigma). Neurons were held at -65 mV and the access resistance of the perforated cells was monitored constantly before and during recordings by a 5 mV seal test. An access resistance of 50 MΩ was considered acceptable to start recording. Recordings were excluded if the resting membrane potential exceeded -50 mV or if the series resistance after the recording deviated more than 30% from its original value. GABA_A_ currents were induced by upon local application of 50 μM muscimol dissolved in HEPES-ACSF (containing in mM: 135 NaCl, 3 KCl, 2.5 CaCl_2_, 1.3 MgCl_2_, 1.25 Na_2_H_2_PO_4_, 20 Glucose, and 10 HEPES) every 30 sec using a Picospritzer II (20 ms). Muscimol responses were recorded in the presence of 1 μM TTX (Abcam) at holding potentials between -100 mV and -30 mV in 10 mV steps. We plotted response amplitude as a function of holding potential and calculated the chloride reversal potential from the intersection of the linear current-voltage curve with the x-axis. To assess the size of GABA_A_ currents independent of holding and reversal potential, we calculated the slope of the current-voltage-plot.

For recordings of spontaneous inhibitory postsynaptic currents (sIPSCs) and miniature inhibitory currents (mIPSCs) we performed whole-cell patch clamp recordings from CA1 pyramidal neurons using borosilicate glass pipettes with resistances of 3-6 MΩ. Pipettes were filled with a KCl-based internal solution (containing in mM: 70 K-gluconate, 70 KCl, 0.5 ethyleneglycol-bis(β-aminoethyl)-N,N,Nʹ,Nʹ-tetraacetic Acid (EGTA), 4 Na_2_phosphocreatine, 4 MgATP, 0.4 NaGTP and 10 HEPES; pH adjusted to 7.2 with KOH) and ACSF was supplemented with 20 μM DNQX (Tocris) and 50 μM DL-APV (Tocris). For recordings of miniature IPSCs (mIPSCs), 1 μM TTX (Abcam) was also added to the ACSF. sIPSCs and mIPSCs were recorded for 6 minutes. For wash in experiments, an sIPSC baseline was recorded for 6 minutes and compared to sIPSCs after 20 minutes wash in of 1 μM TTX (Abcam), 0.2 μM K252a (Tocris), 5 μM AM251 (Sigma) or 1 μM agatoxin-IVA (smartox-biotech), again for 6 minutes. Membrane capacitance and series resistance were monitored during the recordings by a 5 mV seal test. Cells were discarded if series resistance was above 35 MΩ or if the resting membrane potential exceeded -50 mV or if the series resistance after the recording deviated more than 30% from its original value.

All data were blinded before analysis. Events were selected using a template in Clampfit10 software. Further analysis was performed using custom-written MATLAB scripts. Rise time of sIPSCs was determined as the time between 10% and 90% of the peak value. The decay tau was fitted with a single exponential function. Only events with a goodness of fit R^2^ ≥ 0.75 were included.

### Two-photon SuperClomeleon imaging and analysis

We performed two-photon chloride imaging in CA1 pyramidal cells using the SuperClomeleon sensor (Grimley et al., 2013) in cultured slices from SClm mice (Herstel et al., 2022). The SClm sensor consists of Cerulean (a CFP mutant) and Topaz (a YFP mutant) joined by a flexible linker. In the absence of chloride, Fluorescence Resonance Energy Transfer (FRET) occurs from the CFP donor to the YFP acceptor. Chloride binding to YFP increases the distance between the fluorophores and decreases FRET (Grimley et al., 2013).

At the recording day, slices were transferred to a recording chamber and continuously perfused with carbonated (95% O_2_, 5% CO_2_) ACSF supplemented with 1 mM Trolox, at a rate of approximately 1 ml/min. Bath temperature was monitored and maintained at 30-32 °C. Two-photon imaging of CA1 pyramidal neurons or VGAT-positive interneurons in the CA1 area of the hippocampus was performed using a customized two-photon laser scanning microscope (Femto3D-RC, Femtonics, Budapest, Hungary) with a Ti-Sapphire femtosecond pulsed laser (MaiTai HP, Spectra-Physics) and a 60x water immersion objective (Nikon NIR Apochromat; NA 1.0) The CFP donor was excited at 840 nm. The emission light was split using a dichroic beam splitter at 505 nm and detected using two GaAsP photomultiplier tubes. We collected fluorescence emission of Cerulean/CFP (485 ± 15 nm) and Topaz/YFP (535 ± 15 nm) in parallel. Of each slice from SClm mice, 2-5 image stacks were acquired in different fields of view. The resolution was 8.1 pixels/μm (1024x1024 pixels, 126x126 μm) with 1 μm z-steps.

Image analysis was performed using ImageJ software, as described before (Herstel et al., 2022). We manually determined regions of interest (ROIs) around individual neuron somata. To select a representative cell population in SClm slices, in each image z-stack we selected four z-planes at comparable depths in which three pyramidal cells were identified that varied in CFP brightness (bright, middle and dark). We subtracted the mean fluorescence intensity of the background in the same image plane before calculating the fluorescence ratio of CFP and YFP. We limited our analysis to cells which were located within 450 pixels from the center of the image, as FRET ratios showed slight aberrations at the edge of our images. We excluded cells with a FRET ratio < 0.5 or > 1.6 to avoid unhealthy cells.

### Protein extraction and Western Blot analysis

Organotypic hippocampal cultures were washed in cold PBS and subsequently lysed in cold protein extraction buffer containing: 150 mM NaCl, 10 mM EDTA 10 mM HEPES, 1% Triton-X100 and 1x protease and 1x phosphatase inhibitors cocktails (Complete Mini EDTA-free and phosSTOP, Roche). Lysates were cleared of debris by centrifugation (14,000 rpm, 1 min, 4°C) and measured for protein concentration before storage at −20°C until use. Lysates were denatured by adding loading buffer and heating to 95°C for 5 min. For each sample an equal mass of proteins was resolved on 4–15 % polyacrylamide gel (Bio-Rad). The proteins were then transferred (300 mA, 3h) onto ethanol-activated Immobilon-P PVDF membrane (Millipore) before blocking with 3% Bovine Serum Albumin in Tris-buffered saline-Tween (TBST, 20 mM Tris, 150 mM NaCl, 0.1% Tween-20) for 1 h. Primary antibodies used in this study were: mouse anti-NKCC1 (T4, Developmental Studies Hybridoma Bank, 1:1000), rabbit anti-KCC2 (07-432, Merck, 1:1000) and s940-pKCC2 (p1551-940, LuBioSciences, 1:1000), mouse anti-tubulin (T-5168, Sigma, 1:10000). Primary antibodies were diluted in blocking buffer and incubated with the blots overnight at 4 °C under gentle rotation. The membrane was washed 3 times 15 minutes with TBST before a 1h incubation of horseradish peroxidase (HRP)-conjugated antibodies (P0447 goat anti-mouse IgG HRP, Dako, 1:2500 or P0399 swine anti-rabbit IgG HRP, Dako, 1:2500), and washed 3 times 15 minutes in TBST again before chemiluminescence detection. For chemiluminescence detection, blots were incubated with Enhanced luminol-based Chemiluminescence substrate (Promega), and the exposure was captured using the ImageQuant 800 system (Amersham). Images were analyzed in ImageJ, by drawing rectangular boxes around each band and measuring average intensities. Protein levels were normalized to the Tubulin loading controls.

### Immunohistochemistry, confocal microscopy and analysis

Slices were fixed 4 % paraformaldehyde solution in PBS for 30 minutes at room temperature. After washing slices in phosphate buffered saline (PBS), they were permeabilized for 15 minutes in 0.5 % Triton X-100 in PBS for 15 minutes, followed by 1 hour in a blocking solution consisting of 10 % normal goat serum and 0.2 % Triton X-100 in PBS. Slices were incubated in primary antibody solution at 4 °C overnight. The following primary antibodies were used: rabbit polyclonal anti-KCC2 (07-432, Merck, 1:1000), guinea pig anti-NeuN (ABN90, Merck, 1:1000), chicken anti-VGAT (131 006, Synaptic Systems, 1:1000), Rabbit anti-PV (PA1-933, Life Technologies, 1:1000), Mouse anti-CB1R (258 011, Synaptic Systems, 1:1000). Slices were washed in PBS and incubated in secondary antibody solution for four hours at room temperature. Secondary antibodies used were goat anti-rabbit Alexa fluor 647 (A21245, Life Technologies, 1:500), goat anti-guinea pig Alexa Fluor 568 (A11075, Life Technologies, 1:500), goat anti-chicken Alexa fluor 647 (A21449, Life technologies, 1:500), goat anti-rabbit Alexa Fluor 405 (A31556, Life Technologies, 1:500), Goat anti-mouse Alexa fluor 488 (A11029, Life Technologies, 1:500). After another PBS wash, slices were mounted in Vectashield mounting medium (Vector labs).

Confocal images were taken on a Zeiss LSM-700 confocal laser scanning microscopy system with a Plan-Apochromat 40x 1.3 NA immersion objective for KCC2 staining and 63x 1.4 NA oil immersion objective for inhibitory synapse staining. KCC2 staining was imaged in two fields of view in the CA1 *sPyr* of each slice. Inhibitory synapses were imaged in the CA1 *sPyr* and *sRad* of each slice. Image stacks (1024x1024 pixels) were acquired at 6.55 pixels/μm for KCC2 staining, 10.08 pixels/μm for VGLUT/VGAT staining. Step size in z was 0.5 μm for KCC2 staining and 0.4 μm for inhibitory synapse staining.

Confocal images were blinded before analysis. For the analysis of KCC2 levels in control and furosemide-treated slices, ROIs were drawn around 5 NeuN positive cell bodies and this was repeated in 1-3 z-planes (depending on the depth of KCC2 signal) at comparable depths, to measure average KCC2 fluorescence of 5-15 neurons per image stack. For analysis of inhibitory synapses a custom-made macro was used (Ruiter et al., 2020). Briefly, three Z-planes were averaged and background was subtracted with rolling ball radius of 10 pixels. VGAT puncta were identified using watershed segmentation. PV and CB1R channels were thresholded and colocalization with VGAT puncta was analyzed.

### Statistical Analyses

Statistical analysis was performed in Prism (Graphpad). Normality was tested using the D’Agostino & Pearson test. For the comparison of two groups we either used an unpaired Student’s t test (UT; parametric), a Mann-Whitney test (MW; non-parametric), a paired Student’s t test (PT; parametric) or a Wilcoxon signed-rank test (WSR; non-parametric). For comparison of multiple groups, a Kruskal Wallis test (KW; nonparametric) was used, followed by a Dunn’s Multiple Comparison posthoc test (DMC) or a two way ANOVA (2W ANOVA; parametric), followed by a by a Sidak’s Multiple Comparisons posthoc test (SMC). Data are presented as mean ± standard error of the mean. Significance is reported as *p<0.05; **p≤0.01; ***p≤0.001.

## Results

### Furosemide depolarizes GABA responses acutely, but hyperpolarizes GABA responses after one week treatment

We used perforated patch recordings upon muscimol application in CA1 pyramidal neurons of cultured hippocampal slices to record the reversal potential of GABA_A_ receptor currents. Acute administration of 200 μM furosemide in DIV2 slices did not affect the GABA reversal potential (E_GABA_) (Fig. 1A) or GABA driving force (GABA DF) (Fig. 1B), in agreement with a low expression of KCC2 at this age (Salmon et al., 2020; Peerboom et al., 2023). At DIV21, when KCC2 expression is high (Salmon et al., 2020; Peerboom et al., 2023), acute furosemide application elevated E_GABA_ and GABA DF (Fig. 1C,D). This is in agreement with previous reports (Thompson and Gahwiler, 1989; Jarolimek et al., 1999; Deeb et al., 2013; Wright et al., 2017) and comparable with acute inhibition of KCC2 by the specific blocker VU0463271 (VU) in slices at the same developmental stage (Peerboom et al., 2023).

**Figure 1.**
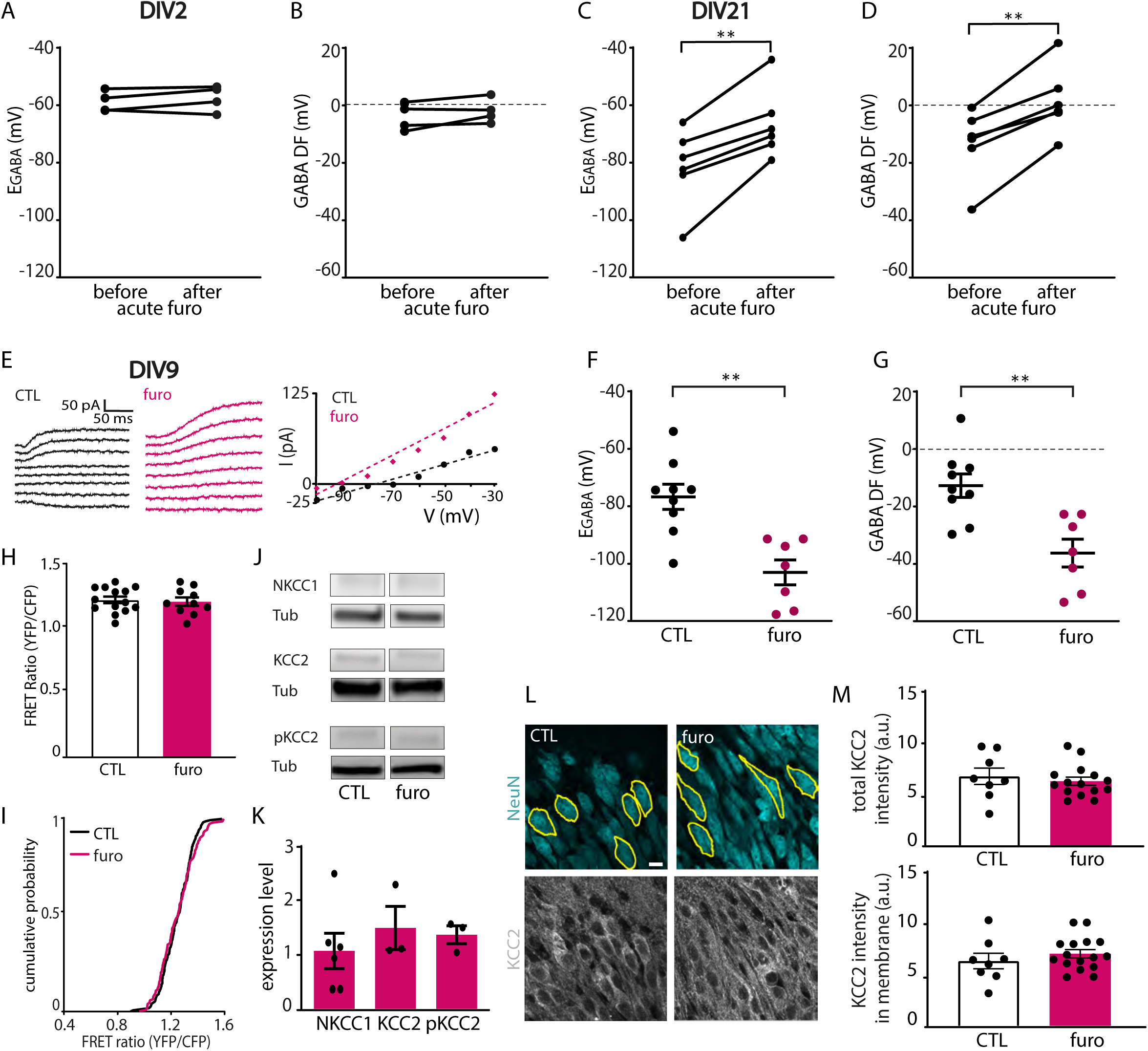
Furosemide acutely depolarizes the GABAergic driving force, but results in hyperpolarization after one week treatment. A, B) GABA reversal potential (EGABA) (PT, p=0.33) and GABA Driving Force (GABA DF) (PT, p=0.19) values recorded in CA1 pyramidal cells at DIV2 before and after acute application of furosemide. Data from 4 cells, 4 slices and 4 mice. C, D) EGABA (MW, p=0.004) and GABA DF (MW, p=0.002) values recorded in CA1 pyramidal cells at DIV21 before and after acute application of furosemide. Data from 6 cells, 6 slices and 5 mice. E) Perforated patch-clamp recording in CA1 pyramidal cells in furosemide-treated and control slices at DIV9. Responses to muscimol application were recorded at holding potentials from -100 mV to -30 mV with 10 mV increments. The GABA reversal potential was determined from the intersection of the linear current-voltage curve with the x-axis. F,G) EGABA (MW, p=0.001) and GABA DF (MW, p=0.005) recorded in CA1 pyramidal cells at DIV9 in control (CTL) and furosemide-treated slices. Data from 7-9 cells, 5-8 slices and 4-5 mice per group. H) Average SClm FRET ratios in CTL and furosemide-treated slice cultures (UT, p=0.75). Data from 10-14 slices and 6-10 mice per group. I) Cumulative distribution of FRET ratios in individual cells in CTL and furosemide -treated cultures. (KS, p=0.23). Data from 15 randomly selected cells per slice. J) Lower panel: Western blots of NKCC1, KCC2 and S940-pKCC2 (pKCC2) protein levels in CTL and furosemide-treated slice cultures. Tubulin (Tub) was used as loading control. K) Summary of data for NKCC1, KCC2 and for s940-pKCC2 (pKCC2) protein levels. Values were normalized to the protein level in CTL cultures (NKCC1: MW, p>0.99; KCC2: MW, p=0.70; pKCC2: MW, p=0.48). Data from 3-5 experiments and 3 mice per group. L) Confocal images of NeuN and KCC2 staining in CTL and furosemide-treated cultures. Yellow lines indicate the outlines of individual cells. Scale bar: 10 μm. M) Total KCC2 levels (UT, p=0.55) and KCC2 levels in membrane (UT, p=0.37) in CTL and furosemide-treated cultures. Each datapoint represents the mean KCC2 intensity of all neurons in one image. Data from 8-17 images, 4-8 slices and 3 mice per group.

We then treated cultured hippocampal slices from DIV1 to DIV8 with 200 μM furosemide. Surprisingly, we observed a strong hyperpolarization of E_GABA_ and a negative shift of the GABA DF at DIV9 after one week of furosemide treatment (Fig. 1E-G). Please note that furosemide was not present during the recordings. Since furosemide has been shown to inhibit α6- and α4-subunit containing GABA_A_ receptors (Thompson and Gahwiler, 1989; Pearce, 1993; Korpi et al., 1995), we checked if GABA_A_ receptor inhibition may have interfered with our measurements of E_GABA_ and GABA DF. As a rough indication, we determined the slope of the current-voltage (IV) plot to assess GABA response amplitudes independently of holding potential and reversal potential. However, IV-slopes were not different (MW, p=0.46), suggesting that furosemide did not have a major effect on GABA_A_ receptors.

A hyperpolarizing shift in E_GABA_ suggest that furosemide treatment resulted in a reduction of intraneuronal chloride or HCO_3_^−^ levels. To directly assess intracellular chloride levels after furosemide treatment, we employed the chloride sensor SuperClomeleon (SClm) (Grimley et al., 2013; Boffi et al., 2018; Rahmati et al., 2021). We used slices from SClm mice in which CA1 pyramidal cells express the SClm sensor and we recorded FRET ratios in control and furosemide-treated slices using two-photon microscopy (Herstel et al., 2022). Remarkably, we did not observe a difference in SClm FRET ratios after one week of furosemide treatment (Fig. 1H,I). The discrepancy between our results from perforated patch recordings and SClm imaging may be due to the decreased sensitivity of the SClm sensor for low chloride levels (Grimley et al., 2013; Herstel et al., 2022). Alternatively, it may reflect concomitant changes in chloride and HCO_3_^−^, as SClm displays substantial pH-sensitivity (Grimley et al., 2013; Boffi et al., 2018; Lodovichi et al., 2022). These possibilities are discussed in more detail in the discussion section.

To examine if the hyperpolarizing GABA signaling after furosemide treatment was due to changes in chloride cotransporter expression, we assessed total NKCC1, KCC2 protein levels in control and furosemide-treated slices using Western blots. We also assessed phosphorylated KCC2 at serine 940 (S940-pKCC2), as this phosphorylation regulates KCC2 function during development (Lee et al., 2007; Mòdol et al., 2014). No significant changes in the levels of NKCC1, KCC2 or S940-pKCC2 were observed (Fig. 1J,K). In addition, we examined KCC2 localization in treated and control slices using immunohistochemistry. We used NeuN to identify individual cell bodies and we estimated KCC2 levels in somata and membranes. We previously showed that we can detect a reduction in KCC2 levels after shRNA using these methods (Peerboom et al., 2023). Furosemide treatment did not significantly affect KCC2 levels in somata or membranes of CA1 pyramidal neurons (Fig. 1L,M). Together, this indicates that furosemide treatment during DIV1 to DIV8 hyperpolarized GABA signaling without changing the expression or surface levels of KCC2, nor by changing expression levels of NKCC1 and S940-pKCC2. These results suggest that furosemide treatment hyperpolarized E_GABA_ via inhibition of chloride transporters KCC2 and NKCC1 without altering their expression.

### Furosemide treatment does not affect inhibitory synapses at DIV9

The results above show that one week of furosemide treatment resulted in a strong hyperpolarization of E_GABA_. Next, we assessed the consequences of this altered GABA signaling on inhibitory transmission. We recorded spontaneous and miniature inhibitory currents (sIPSCs and mIPSCs) using whole-cell voltage clamp recordings in CA1 pyramidal cells in control and furosemide-treated slices at DIV9. We found that the frequency and amplitude of sIPSCs were similar in control and furosemide-treated slices at DIV9 (Fig. 2A,B). There were also no differences in rise and decay kinetics and the membrane capacitance (C_m_) of CA1 pyramidal neurons (Fig. 2C-E). This indicates that GABA signaling does not directly affect the development of inhibitory synapses at this developmental stage (Peerboom et al., 2023).

**Figure 2.**
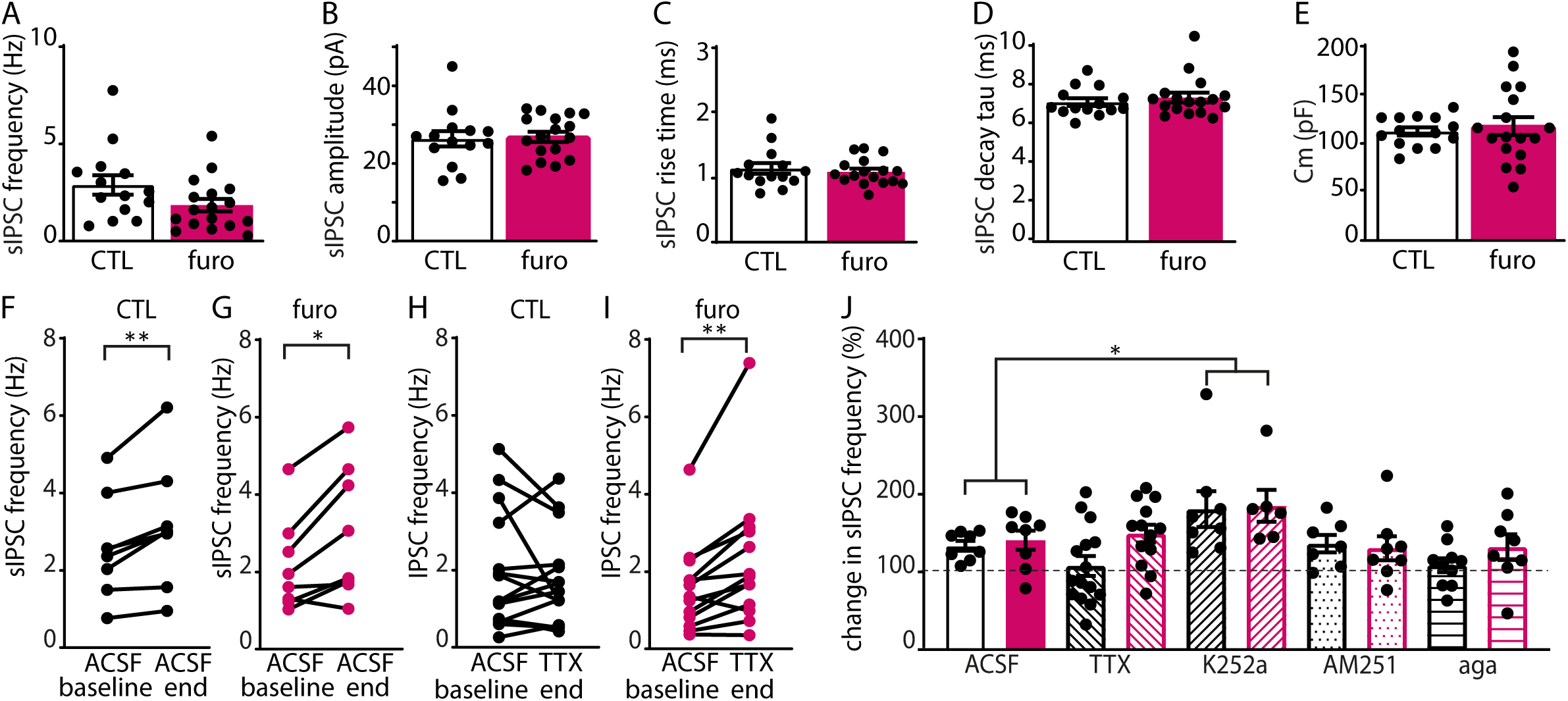
Furosemide does not affect inhibitory transmission at DIV9. A-E) sIPSC frequency (MW, p=0.084), amplitude (UT, p=0.83), risetime (MW, p=0.80) and decay tau (MW, p=0.57) and membrane capacitance (Cm) (UT, p=0.63) in control (CTL) and furosemide-treated slice cultures at DIV9. Data from 14-17 cells, 9 slices and 9 mice per group. F-I) sIPSC frequency after ACSF wash in as a control CTL (WSR, p=0.008) and furosemide-treated (WSR, p=0.002) slice cultures at DIV9. sIPSC frequency after TTX wash in of in CTL (WSR, p=0.80) and furosemide-treated WSR, p=0.002) slice cultures at DIV9. Data from 8-15 cells, 8-15 slices and 7-10 mice per group. J) Change in sIPSC frequency after ACSF wash in as a control, versus after wash in of TTX, K252a, AM251 and agatoxin-IVA (aga) to block neuronal activity, BDNF, endocannabinoids and P/Q calcium channels respectively in control and furosemide-treated slices at DIV9 (2W ANOVA, treatment p=0.18, blocker p=0.002; SMC, control versus TTX, p=0.95; control versus K252a, p=0.020; control versus AM251, p=0.99; control versus agatoxin-IVA (aga), p=0.82). Data from 7-15 cells, 7-15 slices and 4-10 mice per group.

When recording sIPSCs, we noticed that the sIPSC frequency slightly increased by ∼20% during the first 20 minutes of recording in both control and furosemide-treated slices (Fig. 2F,G), which was associated with a gradual increase of the fraction of sIPSCs with small amplitudes and slow rise times (data not shown). This suggests that the detection of sIPSCs from distal locations gradually improved over time due to diffusion of high chloride internal solution from the recording pipette into the recorded cell. When we washed in TTX, this ‘run-up’ was compensated by the simultaneous reduction of activity-dependent IPSCs in control slices (Fig. 2H). However, in furosemide-treated slices the frequency increase remained during wash-in of TTX (Fig. 2I), suggesting that a smaller fraction of sIPSCs were activity-dependent. We assessed if this difference was due to increased levels of activity-dependent factors that suppress GABA release in furosemide-treated slices, such as endocannabinoids or BDNF. We assessed the contribution of the BDNF receptor TrkB by washing in TrkB antagonist K252a during sIPSC recordings in control and furosemide-treated slices. We found that sIPSC frequency was enhanced after wash-in of K252a, suggesting that there is ongoing suppression of GABA release by BDNF in our slices at DIV9. However, there was no differential contribution of BDNF between control and furosemide-treated slices (Fig. 2J). We also washed in AM251, a specific antagonist of the endocannabinoid receptor CB1. We found that endocannabinoid signaling did not have a prominent influence on sIPSCs, and there was no difference between control and furosemide-treated slices (Fig. 2J). Blocking presynaptic P/Q calcium channels with agatoxin-IVA (aga) (Poncer et al., 1997; Goswami et al., 2012) also did not significantly alter sIPSC frequency in control and furosemide-treated slices (Fig. 2J). These data cannot explain an increase in relative contribution of activity-independent release in sIPSCs after furosemide treatment at DIV9.

### Inhibitory transmission is increased in furosemide-treated slices at DIV21

Two weeks after ending the treatment at DIV21, E_GABA_ and GABA DF in pyramidal neurons had normalized and reached control levels (Fig. 3A,B). We previously found that a transient elevation of chloride using VU treatment from DIV1 to DIV8 resulted in indirect changes in inhibitory transmission at DIV21 (Peerboom et al., 2023). We therefore assessed if furosemide treatment from DIV1 to DIV8 affected sIPSCs at DIV21 (Fig. 3C). There was large variability in sIPSC frequency between cells, but we did not detect any significant difference between furosemide-treated and control slices (Fig. 3D). We did observe a small increase in sIPSC amplitude in furosemide-treated slices (Fig. 3E), while the rise and decay time of the sIPSCs were not different (Fig. 3F, G). We noticed that during these sIPSC recordings the cell capacitance C_m_ of CA1 pyramidal cells gradually decreased in furosemide-treated slices, while C_m_ values were stable in control slices (Fig. 3H). As resting membrane potential and input resistance of all recorded cells remained stable, this decrease in C_m_ appears to be induced by exchange of the cytoplasm of the cells with the high chloride internal solution. We speculate that it may reflect an impaired volume regulation upon chloride load (see discussion). To assess whether the changes in inhibitory transmission were activity-dependent, we also recorded mIPSCs (Fig. 3I). We observed that mIPSCs frequency was increased in furosemide-treated slices at DIV21, while amplitude, rise and decay time were not different from control slices (Fig. 3J-M). For the mIPSC recordings we waited at least 5 minutes before starting the recordings and whole-cell capacitance was stable and not different in control and furosemide-treated slices (Fig. 3N). The observed increase in mIPSC frequency without an accompanying change in sIPSC frequency indicates that activity-independent GABA release was specifically increased in furosemide-treated slices at DIV 21.

**Figure 3.**
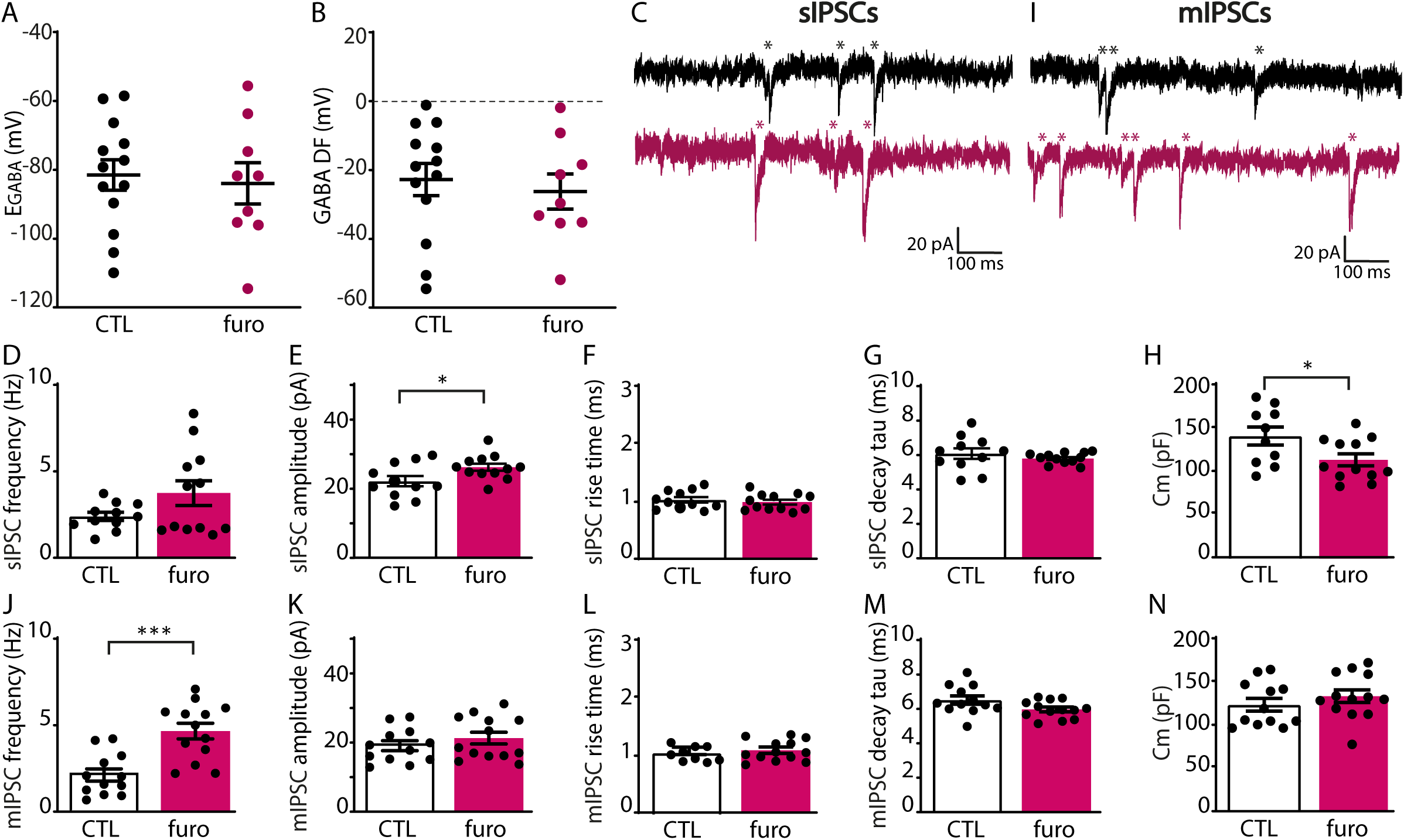
Furosemide increases inhibitory transmission at DIV21. A,B) GABA reversal potential (EGABA) (MW, p=0.56) and GABA Driving Force (GABA DF) (MW, p=0.76) in control (CTL) and furosemide-treated slice cultures at DIV21. Data from 9-13 cells, 9 slices and 9 mice per group. C) sIPSC recording from CA1 pyramidal cells in CTL and furosemide-treated organotypic cultures at DIV21. sIPSCs are indicated with *. D-G) sIPSC frequency (UT, p=0.10), amplitude (UT, p=0.034), risetime (UT, p=0.54) and decay tau (UT, p=0.34) in CTL and furosemide-treated slice cultures at DIV21. I) Membrane capacitance (Cm) of CA1 pyramidal cells in CTL and furosemide-treated slice cultures at DIV21 (UT, p=0.035). I) mIPSC recording from CA1 pyramidal cells in control and furosemide-treated organotypic cultures at DIV21. mIPSCs are indicated with *. J-M) mIPSC frequency (UT, p=0.0002), amplitude (UT, p=0.34), risetime (MW, p=0.73) and decay tau (UT, p=0.06) in CTL and furosemide-treated organotypic cultures at DIV21. N) Membrane capacitance (Cm) of CA1 pyramidal cells in the presence of TTX in CTL and furosemide-treated slice cultures at DIV21 (UT, p=0.35). Data in D-H from 11-12 cells, 5-6 slices and 3-5 mice per group; data in J-N from 12-13 cells, 5 slices and 4 mice per group.

A change in mIPSC frequency while the amplitudes remain unaffected could reflect an increase in the number of inhibitory synapses in furosemide-treated slices compared to control slices at DIV21. To assess synapse numbers, we used immunohistochemistry to visualize inhibitory synapses in *stratum Pyramidale (sPyr*) and *stratum Radiatum* (*sRad*) using the vesicular GABA transporter VGAT. In addition, we stained for CB1 receptors (CB1R) and parvalbumin (PV), to identify inhibitory synapses from specific interneuron subtypes (Fig. 4A,B). The density and size of VGAT puncta in the CA1 area were similar in control and furosemide-treated slices at DIV9 (Fig. 4C-F) and we did not observe any differences in the density of CB1R+ and PV+ VGAT puncta (Fig. 4G-I). At DIV21 however (Fig. 5A,B), we observed an increased density and reduced size of VGAT puncta in the *sRad* in furosemide-treated slices compared to controls (Fig. 5E,F), but not in the *sPyr* (Fig. 5C,D). The density of PV- and CB1R-positive puncta were not different in furosemide-treated and control slices (Fig. 5G-I), suggesting that the increase in VGAT puncta is due to synapses from other interneuron subtypes. To assess possible changes in cell size, we analyzed soma area from the NeuN staining, but we found no differences between furosemide-treated and control slices (control: 211 ± 7 μm^2^ and furosemide: 211 ± 13 μm^2^; p=0.9757, UT-test; 14-16 images from 7-8 slices from 3 mice). These results suggest that the density of inhibitory synapses in the dendritic region was increased at DIV21 in furosemide-treated slices.

**Figure 4.**
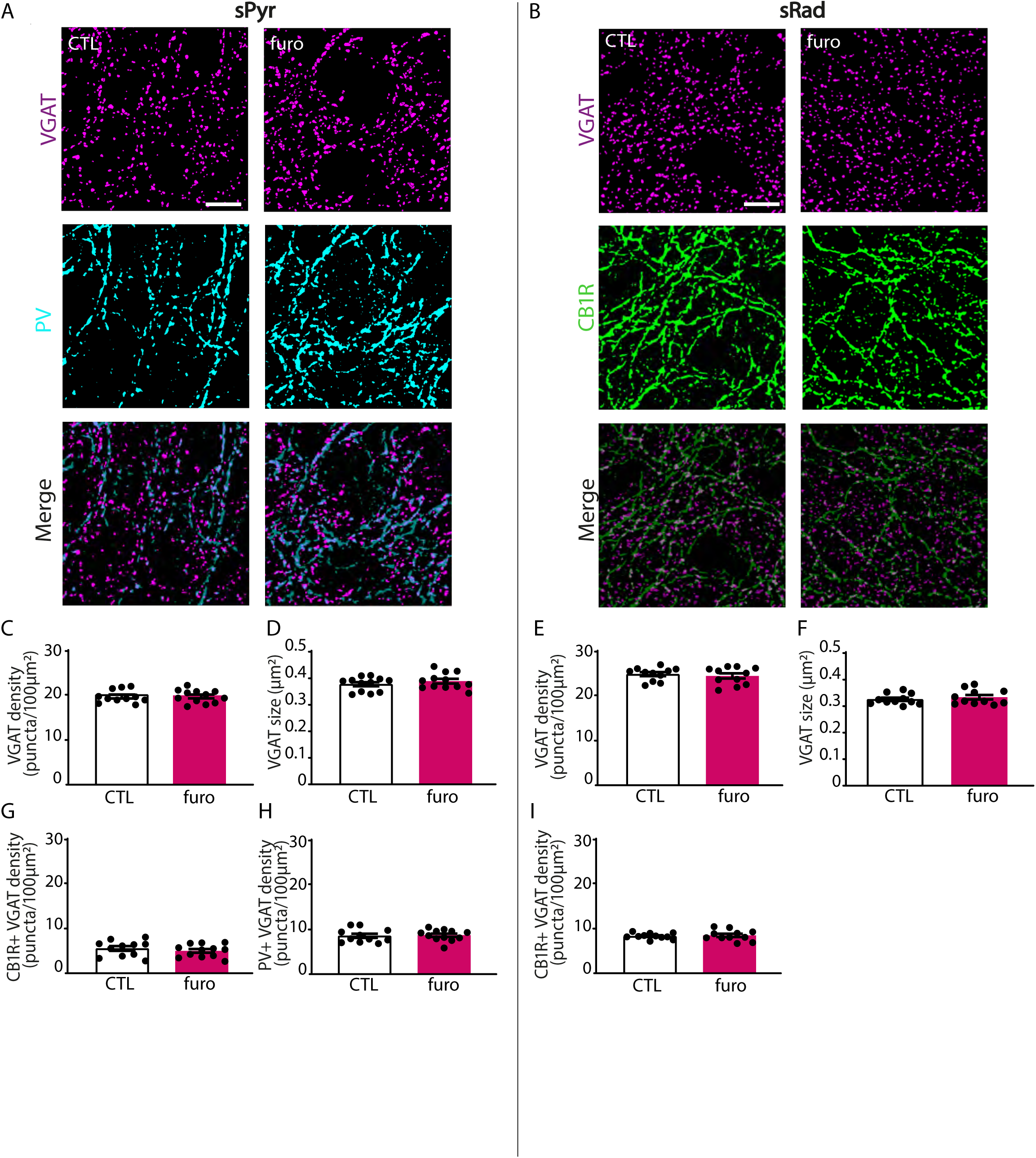
Furosemide does not affect inhibitory synapses at DIV9. A,B) VGAT, PV and CB1R immunofluorescence in the CA1 *stratum Pyramidale* (*sPyr)* and *stratum Radiatum* (*sRad)* of control (CTL) and furosemide-treated organotypic cultures at DIV9. Scale bar=10 μm. C,D) The density (UT, p=0.50) and size (UT, p=0.47) of VGAT puncta in the *sPyr* of CTL and furosemide-treated organotypic cultures at DIV9. E,F) The density (UT, p=0.59) and size (UT, p=0.46) of VGAT puncta in *sRad* of CTL and furosemide-treated organotypic cultures at DIV9. G,H) The density of CB1R-positive VGAT puncta (UT, p=0.43) and PV positive VGAT puncta (UT, p=0.82) in *sPyr* of CTL and furosemide-treated organotypic cultures at DIV9. I) The density of CB1R-positive VGAT puncta (UT, p=0.68) in *sRad* of CTL and furosemide-treated organotypic cultures at DIV9. Data in C-I from 11-12 images, 6 slices and 2 mice per group.

**Figure 5.**
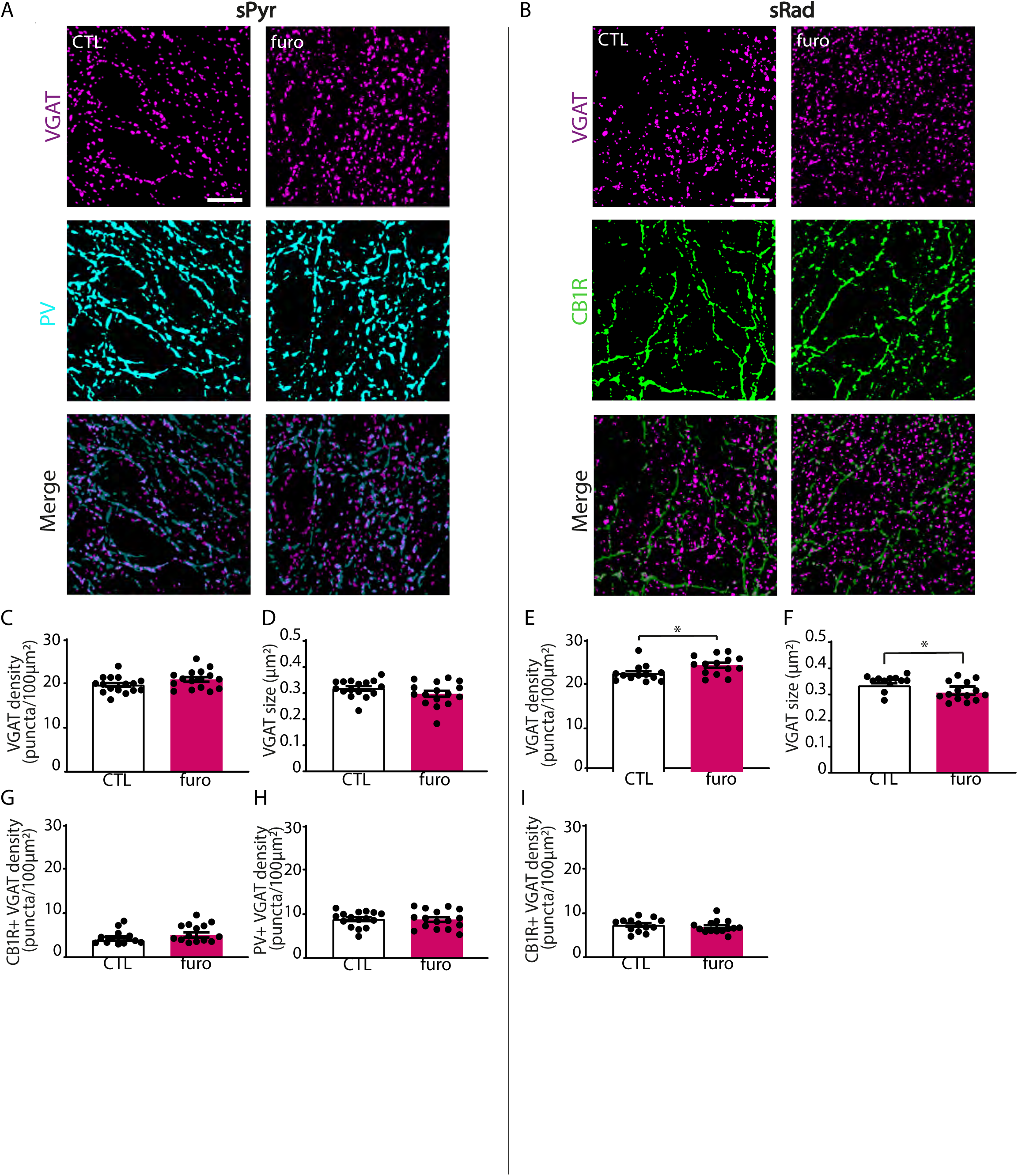
Furosemide increases the number of VGAT puncta at DIV21 in sRad. A,B) VGAT, PV and CB1R immunofluorescence in the CA1 *stratum Pyramidale* (*sPyr)* and *stratum Radiatum* (*sRad)* of control (CTL) and furosemide-treated organotypic cultures at DIV21. Scale bar=10 μm. C,D) The density (UT, p=0.09) and size (UT, p=0.14) of VGAT puncta in *sPyr* of CTL and furosemide-treated organotypic cultures at DIV21. Data from 16 images, 8 slices and 3 mice per group. E,F) The density (MW, p=0.019) and size (MW, p=0.015) of VGAT puncta in *SRad* of CTL and furosemide-treated organotypic cultures at DIV21. Data from 13 images, 7 slices and 3 mice per group. G,H) The density of CB1R (UT, p=0.11) and PV positive VGAT puncta (UT, p=0.89) in *sPyr* of CTL and furosemide-treated organotypic cultures at DIV21. Data from 16 images, 8 slices and 3 mice per group. I) The density of CB1R-positive VGAT puncta (MW, p=0.30) in *SRad* of CTL and furosemide-treated organotypic cultures at DIV21. Data from 13 images, 7 slices and 3 mice per group.

## Discussion

In this study, we used furosemide to manipulate the GABA shift in organotypic hippocampal slices. Unexpectedly, we observed that treatment with furosemide from DIV1 to DIV8 had an opposite effect compared to acute furosemide application. While acute application depolarized the reversal potential for GABA signaling (E_GABA_), one week furosemide treatment resulted in a strong hyperpolarizing shift of E_GABA_. This shift was not accompanied by a change in SuperClomeleon FRET ratios or chloride co-transporter expression. We observed that sIPSCs were not affected by the hyperpolarization of E_GABA_ after furosemide treatment at DIV9, consistent with the notion that GABA signaling in the second postnatal week no longer promotes synapse formation (Wang and Kriegstein, 2011; Salmon et al., 2020; Peerboom et al., 2023). However, we found that at DIV21, two weeks after ending furosemide treatment and when E_GABA_ had normalized, inhibitory currents were increased. This was accompanied by an increase of the density of VGAT puncta in *sRad*. Together, these results show that accelerating the GABA shift with furosemide does not directly affect the development of inhibitory synapses. However, transient furosemide treatment appeared to indirectly alter increase inhibitory transmission two weeks later. Unfortunately, our data do not provide a clear mechanism explaining these observations, but we offer our thoughts and speculations below.

### Effect of furosemide on neuronal chloride levels

The depolarizing effect of acute furosemide application at DIV21 was comparable to the depolarization after acute inhibition of KCC2 by specific KCC2 blocker VU0463271 (VU) in slices at the same developmental stage (Peerboom et al., 2023), suggesting that acutely applied furosemide mainly inhibits KCC2. At the concentration used here (200 μM), furosemide is expected to block both KCC2 and NKCC1 (Orlov et al., 2015). However, NKCC1 is mainly inactive in adult tissue and maintains chloride levels only until approximately P15-21 (Romo-Parra et al., 2008; Sulis Sato et al., 2017; Salmon et al., 2020). Previous studies also reported a depolarizing effect of furosemide when acutely applied to (slice) cultures at a developmental stage equal to P14 or later (Thompson and Gahwiler, 1989; Jarolimek et al., 1999; Deeb et al., 2013; Wright et al., 2017). In our slices at DIV21 (prepared from P7 mice), NKCC1 may therefore no longer directly regulate chloride levels, which explains why acute furosemide application only acted via KCC2 at DIV21.

In stark contrast, furosemide treatment from DIV1 to DIV8 resulted in hyperpolarizing E_GABA_. As furosemide blocks both main chloride transporters KCC2 and NKCC1, we had predicted that intracellular chloride levels would roughly follow the Nernst equilibrium. The observed hyperpolarized E_GABA_ suggests that an additional chloride extrusion mechanism exists that remains unidentified here. We checked that furosemide treatment did not affect expression levels of chloride transporters. If furosemide treatment hyperpolarized E_GABA_ by reducing intracellular chloride levels, we would expect to observe an increase in SClm FRET ratios (reflecting a decrease in chloride levels) after furosemide treatment, which was not the case. However, although the SClm sensor is useful to detect increases in chloride levels (Peerboom et al., 2023), its sensitivity of is reduced at chloride concentrations below ∼5 mM (Grimley et al., 2013), which is close to physiological chloride levels in neurons (Boffi et al., 2018). This suggests that chloride levels in our DIV9 slices after furosemide treatment are too low to detect a further decrease with the SClm sensor.

### Possible effect of furosemide on pH

In addition to blocking chloride transporters, furosemide may interfere with enzymes that are involved in HCO_3_^-^ regulation (Halligan et al., 1991; Temperini et al., 2009; Uwera et al., 2015). GABA_A_ receptors are permeable for chloride and HCO_3_^−^ ions with a permeability of around 70% and 30% respectively (Kaila et al., 2014). The reversal potential of HCO_3_^−^ is maintained by pH-regulatory proteins at around -10 mV (Kaila et al., 2014), which is much more positive than the chloride reversal potential E_Cl_). As a result, E_GABA_ is slightly more positive than E_Cl_ and the flow of chloride through GABA_A_ receptors is accompanied by an outflow of HCO_3_^−^ (Kaila et al., 2014). The lack of change in SClm FRET ratios after furosemide treatment raises the possibility that the observed hyperpolarization of E_GABA_ reflect changes in intracellular HCO_3_^−^ levels, rather than chloride. If furosemide would decrease intraneuronal HCO_3_^−^ levels, the outflow of HCO_3_^−^ upon GABA_A_ receptor activation would be reduced, and E_GABA_ would become more negative. Previous studies have suggested that furosemide can inhibit the Cl^−^/HCO_3_^−^ anion exchanger AE3 (Halligan et al., 1991; Uwera et al., 2015). AE3 exchanges intracellular HCO_3_^−^ for extracellular chloride, thereby lowering intracellular pH and increasing intracellular chloride concentrations in response to intracellular alkali loads (Gonzalez-Islas et al., 2009; Pfeffer et al., 2009; Romero et al., 2013). Furosemide can also block carbonic anhydrase (CA) (Temperini et al., 2009), which is a cytosolic enzyme that mediates the conversion of HCO_3_^−^ (CO_2_ + H_2_O ⇄ HCO_3_^−^+ H^+^) to buffer intraneuronal pH after alkali or acid loads (Ruusuvuori and Kaila, 2014). To our knowledge there are no studies directly examining the contribution of AE3 or CA to E_GABA_. Theoretically, inhibition of AE3 by furosemide would decrease intracellular chloride levels while increasing HCO_3_^-^, which have opposing effects on E_GABA_. Inhibition of CA would in theory reduce the contribution of E_HCO3_ to E_GABA_ and result in a negative shift of E_GABA_ towards E_Cl_. If the hyperpolarization of E_GABA_ that we observed in our experiments were solely due to changing HCO_3_^−^ levels without changing chloride, pH values would be unphysiologically low. As neurons were generally healthy after furosemide treatment, such a large pH change seems unlikely. However, a small reduction in intracellular pH would decrease SClm FRET ratios (Grimley et al., 2013; Boffi et al., 2018; Lodovichi et al., 2022), which may have obscured an increase in FRET ratios due to a decrease in chloride levels. We therefore conclude that furosemide treatment most likely resulted in a decrease in chloride levels, but our data do not exclude an accompanying subtle change in pH regulation.

### Possible effect of furosemide on cell volume regulation

We observed that the membrane capacitance of CA1 pyramidal neurons gradually reduced after break-in for whole-cell recordings in furosemide-treated slices, but not in control slices at DIV21 (Fig. 3H), and also not at DIV9 (Fig. 2E). A reduction in whole cell membrane capacitance may reflect a reduction in cell size (Taylor, 2012). Previous studies have shown that furosemide inhibits activity-induced changes in the volume of neuronal tissue during epilepsy (Hochman et al., 1995; Gutschmidt et al., 1999) via inhibition of NKCC1 on glia cells (Ransom et al., 1985; Hochman, 2012). Glia disperse local, activity-induced elevations in potassium and chloride ions which are followed by water and result in glial swelling. As capacitance was not different immediately after break-through, cell volume did not appear to be affected per se. This was also supported by the observation that soma sizes of CA1 pyramidal cells were found similar in furosemide-treated and control slices. Overall cellular health was not affected in furosemide-treated slices at DIV21 as resting membrane potential and input resistance remained stable during the recordings and were comparable to control slices. Instead, the observed gradual decrease in whole cell capacitance may indicate a loss of cell volume regulation in response to wash-in of the high chloride pipette solution. We speculate that chloride extrusion mechanisms triggered by the high chloride load (Jarolimek et al., 1999) may have instigated loss of cell volume, resulting in the observed C_m_ decrease. This suggests that the transient furosemide treatment between DIV1 and DIV8 led to long-lasting alterations in chloride homeostasis and/or cell volume regulation.

### Furosemide-treatment indirectly increased inhibitory transmission

We observed that sIPSCs were not affected by the hyperpolarization of E_GABA_ at DIV9, immediately after the furosemide treatment (Fig. 2A-E). These results confirm previous studies reporting that synapse formation is mostly independent from GABA signaling after the first postnatal week (Wang and Kriegstein, 2011; Peerboom et al., 2023). More surprisingly, we found that furosemide treatment resulted in an increase in inhibitory transmission at DIV21. The frequency of mIPSCs were strongly increased, while sIPSC frequency was less affected. This points to a relative increase in activity-independent GABA release at DIV21, which was also noticeable, although much less pronounced, at DIV9. This may indicate subtle alterations in inhibitory synapses, but we did not further examine release properties. The observed increase in sIPSC amplitude may reflect a compensatory change. In our immunohistochemistry experiments we observed an increased density of inhibitory synapses in *sRad* in furosemide-treated slices compared to control slices, which may explain the increase in mIPSC frequency. However, one should be cautious in interpreting synaptic density in the two conditions, as furosemide treatment may have induced tissue shrinkage (see above), which would increase apparent synaptic density. To determine synaptic densities independently of tissue shrinkage, the density of synapses along the dendrites (Feng et al., 2021) or per soma (Bijlsma et al., 2022) should be assessed.

### Possible anti-epileptic actions of furosemide

Furosemide is commonly used as a diuretic, but has important anti-epileptic actions. A case-control study on hypertension and epilepsy showed that diuretics including furosemide protected patients with hypertension against an increased risk on seizures in later life (Hesdorffer et al., 1996). Subsequently, the antiepileptic effects of furosemide were demonstrated in several animal models (Hochman et al., 1995; Gutschmidt et al., 1999; Holtkamp et al., 2003; Barbaro et al., 2004; Viitanen et al., 2010; Hochman, 2012; Uwera et al., 2015; Chen et al., 2022). Since then, several epidemiological and experimental studies have reported anticonvulsant actions of furosemide, but because it broad range of activity the mechanism behind furosemide’s anti-epileptic effects remains mostly unclear (Hesdorffer et al., 1996; Staley, 2002; Maa et al., 2011). Although the precise mechanism remains obscure, our results suggest that furosemide treatment indirectly increases inhibitory transmission, which adds to its previously described anticonvulsant action. An increase in activity-independent GABA release may result in an increase in ambient GABA levels, mediating a cellular hyperpolarization and reducing overall excitability.

### Final remarks

In a recent study we treated slice cultures with VU, a specific blocker of the chloride exporter KCC2, to delay the postnatal GABA shift and maintain a depolarizing E_GABA_ until DIV8. Similar to the current results, we also observed an indirect increase in sIPSC frequency at DIV21 cells (Peerboom et al., 2023). However, the mechanism seems different as mIPSC frequency and synapse numbers were not affected in VU-treated slices (Peerboom et al., 2023). In interpreting our results and that of previous studies, it is hard to disentangle the versatile pharmacological profile of furosemide and the many roles for neuronal chloride in regulating cell volume (Jentsch and Pusch, 2018) and possibly cellular excitability (Huang et al., 2012; Seja et al., 2012; Jentsch and Pusch, 2018; Goutierre et al., 2019; Sinha et al., 2022; Peerboom et al., 2023). Our study further underscores the need for a better understanding of the role of chloride as an intracellular messenger in developing neurons.

## Acknowledgements

This research was supported by a TOP grant from ZonMW (#91216021) and by the Nederlandse Organisatie voor Wetenschappelijk Onderzoek (NWO; #OCENW.KLEIN.150). We thank Prof. Thomas Kuner for providing the Cre-dependent SClm construct (Boffi et al., 2018). We thank Prof. Kevin Staley for providing the SuperClomeleon^lox/-^ mouse line, Dr. Stefan Berger for the CamKIIα^Cre/-^ mice and Dr. Henk Karst for sharing these mice with us. We thank Dunya Selemangel for help with SuperClomeleon data analysis and René van Dorland for excellent technical support.

